# The novel variant IBDV induces inflammatory responses via VP3-mediated pyroptosis by directly targeting Caspase-9

**DOI:** 10.64898/2026.01.12.698959

**Authors:** Tao Zhang, Suyan Wang, Xiaole Qi, Lijie Tang, Yongzhen Liu, Guodong Wang, Hangbo Yu, Yulong Zhang, Ningyu Gao, Ying Wang, Wenrui Fan, Ruihan Zhao, Yuntong Chen, Yanping Zhang, Hongyu Cui, Yulu Duan, Yulong Gao

## Abstract

The novel variant infectious bursal disease virus (nVarIBDV), which has been widely prevalent since 2017, threatens the poultry industry by inducing severe bursal atrophy and intense inflammatory responses. Understanding the inflammatory mechanism underlying nVarIBDV infection is critical for preventing virus-induced damage. Pyroptosis, an inflammatory type of programmed cell death, may contribute to the inflammation and bursa of Fabricius damage induced by nVarIBDV. Here, we found that the nVarIBDV infection induced severe inflammatory responses both *in vivo* and *in vitro*, which were associated with pyroptosis. Further research revealed that the viral protein VP3 drove pyroptosis induced by nVarIBDV infection by indirectly activating the Caspase-3-GSDME pathway, leading to GSDME cleavage. Mechanistically, VP3 directly interacted with and activated Caspase-9, thereby initiating Caspase-9-Caspase-3-GSDME pathway. More importantly, the Serine 33 of VP3 was identified as the key amino acid for interacting with and activating Caspase-9. Mutation of this residue significantly weakened the ability of VP3 to interact with Caspase-9 and activate the Caspase-9-mediated pyroptosis pathway, altered the binding mode of the Caspase-9-VP3 protein complex, and ultimately reduced the ability of VP3 to induce pyroptosis. In conclusion, our results elucidated a novel mechanism by which nVarIBDV infection induced inflammatory responses whereby viral protein VP3 triggered pyroptosis by targeting the Caspase-9-Caspase-3-GSDME pathway.

**IMPORTANCE:** nVarIBDV is globally prevalent and poses a significant threat to the poultry industry by causing inflammation and damage to the bursa of Fabricius. This study revealed that nVarIBDV infection induced inflammatory responses and bursa of Fabricius damage, primarily mediated by the viral protein VP3 targeting Caspase-9 to induce pyroptosis. To our knowledge, this is the first report on the interaction between inflammatory responses induced by IBDV infection and pyroptosis. In addition, serine 33 was identified as the critical amino acid required for maintaining the stability of the interaction between VP3 and Caspase-9 and activation of the Caspase-9-Caspase-3-GSDME cascade pyroptosis pathway and further inflammatory responses. Collectively, our findings not only elucidate a novel mechanism by which nVarIBDV induces pyroptosis and further leads to inflammation but also suggest that the identification of this serine site may represent a potential therapeutic target to block nVarIBDV-induced inflammation and bursa of Fabricius damage.

## INTRODUCTION

Infectious bursal disease virus (IBDV) is a highly contagious pathogen that causes an acute and immunosuppressive disease in chickens aged 3 to 6 weeks, leading to substantial economic losses in the poultry industry (1–3). Its genome consists of two segments, A and B; segment A encodes viral proteins VP2, VP3, VP4, and VP5, and segment B encodes the viral RNA-dependent RNA polymerase VP1 (2). IBDV includes two serotypes: serotype I and serotype II (4). Serotype I is classified into classic type (cIBDV), variant type (varIBDV), and very virulent type (vvIBDV) based on antigenicity and virulence (5, 6). In recent years, a novel variant IBDV strain (nVarIBDV) has emerged and has become widely prevalent in China (7), Japan (8), South Korea (9), Malaysia (10) Argentina (11), Egypt (12) and other regions, causing severe clinical symptoms in chickens. Clinical studies have shown that the pathogenic core of nVarIBDV lies in inducing intense inflammatory responses and tissue damage (13). After nVarIBDV infection, significant upregulation of key inflammatory mediators has been widely observed in the bursa of Fabricius, leading to severe inflammatory pathology characterized by inflammatory cell infiltration, mucus exudation and tissue damage (14). Ultimately, this inflammation-driven immune damage not only causes atrophy of the bursa of Fabricius, but also reduces the immune response of the chicken flock to other vaccines, such as Avian influenza virus (AIV) (15) and Newcastle disease virus (NDV) (16), making them prone to secondary infections and significantly exacerbating the actual harm of nVarIBDV in poultry production. Therefore, elucidating the inflammatory mechanism induced by nVarIBDV is crucial for understanding viral pathogenesis and developing targeted intervention strategies.

Viral infections induce inflammation through various pathways, leading to an imbalance between pro-inflammatory and anti-inflammatory factors and thereby exacerbating pathological damage (17). Severe acute respiratory syndrome coronavirus 2 (SARS-CoV-2) infection can activate the ZBP1-RIPK3 pathway to induce the release of interleukin-1β (IL-1β), aggravating damage of lung tissue (18). Pseudorabies virus (PRV) protein UL4 activates the NLRP3 and AIM2 inflammasomes through a SYK/JNK phosphorylation pathway that enhances ASC oligomerization to induce inflammatory responses and tissue damage (19). Therefore, viruses can regulate the upstream pathways of inflammatory factors to precisely induce inflammatory respones. Furthermore, studies have shown that the H7N9 virus can cause a severe cytokine storm by activating Gasdermin (GSDM) family protein to mediate pyroptosis, leading to lung tissue damage (20). This indicates that pyroptosis is also an important pathway for viral infection to induce inflammation and damage. Pyroptosis is a unique and highly inflammatory programmed cell death (PCD) pathway, the core mechanism of which involves GSDM family proteins (GSDMA, GSDMB, GSDMC, GSDMD and GSDME) being cleaved by Caspase family proteins or other proteins with protease functions, generating N-terminal domains that are located on the cell membrane and have pore-forming activity, ultimately leading to the release of cellular contents (IL-1β and IL-18) and triggering an inflammatory responses (21, 22). Some viruses, such as SARS-CoV-2, directly cleave the

Gasdermin protein, while others, such as NDV, indirectly cleave the Gasdermin protein through the host caspase, ultimately leading to cell death and the release of inflammatory factors (23). Therefore, viruses can precisely trigger and amplify inflammatory responses by directly or indirectly regulating host cell death or inflammatory pathways, ultimately leading to inflammatory responses and tissue damage.

Multiple studies have shown that during IBDV infection, obvious inflammatory cell infiltration and inflammatory responses can be seen in the bursa of Fabricius (13, 14, 24, 25), which is similar to the inflammatory responses induced by the H7N9 virus [18]. However, it remains unclear whether nVarIBDV triggers inflammation by inducing pyroptosis and thereby causes bursa of Fabricius damage.

In this study, we infected SPF chickens and DT40 cells with nVarIBDV and detected the characteristics of inflammation and pyroptosis induced by nVarIBDV, as well as the underlying molecular mechanisms. Our results showed that nVarIBDV infection promoted pyroptosis both *in vivo* and *in vitro*, thereby intensifying the inflammatory responses. Moreover, we demonstrated that VP3 of nVarIBDV activated the Caspase-9-Caspase-3-GSDME pathway through direct interaction with Caspase-9, leading to B lymphocyte pyroptosis. Importantly, the Serine 33 of VP3 was identified as the key amino acid responsible for interacting with Caspase-9 and activating Caspase-9-Caspase-3-GSDME pathway. This study reveals the novel molecular mechanism by which nVarIBDV induces an inflammatory storm by inducing B lymphocyte pyroptosis.

## Materials and methods

### Cells and viruses

Human embryonic kidney (HEK-293T) cells and DF-1 cells were cultured in Dulbecco’ s modified Eagle’ s medium (DMEM; Basal Media, L110KJ) supplemented with 10% fetal bovine serum (FBS, Sigma-Aldrich), 100 U/ml penicillin, and 100 µg/ml streptomycin. Chicken lymphoma cells (DT40) were cultured in RPMI-1640 medium (Sigma-Aldrich) supplemented with 10% FBS, 2% chicken serum (Sigma-Aldrich), 1% sodium pyruvate (Sigma-Aldrich), 100 U/ml penicillin, and 100 µg/ml streptomycin, and 0.1% β-mercaptoethanol (Sigma-Aldrich). HEK-293T and DT40 cells were maintained in a humidified incubator at 37℃ with 5% CO_2_. DF-1 cells were cultured in a humidified incubator containing 5% CO_2_ at 38.5°C. The IBDV strain used in this study was a novel variant IBDV strain SHG19 (26), which was isolated and purified by the Avian Immunosuppressive Disease Research Group of Harbin Veterinary Research Institute (HVRI), Chinese Academy of Agricultural Sciences (hereinafter referred to as the "laboratory").

### Plasmids construction and transfection

The genes encoding IBDV structural and non-structural proteins were amplified from the IBDV SHG19 template. After digestion with *EcoR*I and *Cla*I enzymes, Polymerase Chain Reaction (PCR) amplicons were ligated into pCAGGS to generate the HA- VP1, HA-VP2, HA-VP3, HA-VP4 and HA-VP5 vectors. VP3 truncated mutants (HA-VP3 Δ N30, HA-VP3 Δ N100, HA-VP3 Δ C30, HA-VP3 Δ C100, HA-VP3ΔN40, HA-VP3ΔN60, HA-VP3ΔN80, HA-VP3ΔN35, HA-VP3ΔN31, HA-VP3ΔN32, HA-VP3ΔN33 and HA-VP3ΔN34) were constructed by truncating the corresponding coding sequences. Full-length cDNAs of chicken Caspase-3, GSDME and Caspase-9 were amplified from the reverse-transcribed cDNA derived from DF-1 cells. The GSDME cDNA was then inserted into a modified pCAGGS-Flag-N-C vector, the Caspase-9 cDNA was then inserted into a modified pCAGGS-Flag-C vector, while the Caspase-3 cDNA was inserted into modified pCAGGS-Myc-C and pCAGGS-Flag-Cvector. To further demonstrate the interaction site between VP3 and Caspase-9, VP3 Ser33 mutant (Flag-VP3S33A) was constructed by replacing the targeted Serine with Alanine by using site-directed mutagenesis.

HEK-293T cell transfection was conducted using the PolyJet^TM^ (Signagen, SL100688) *in vitro* DNA according to the manufacturer’s instructions.

DF-1 cell transfection was conducted using the TransIT -X2 (Mirusbio, MIR 6000) according to the manufacturer’s instructions.

### Antibodies and reagents

The mouse anti-HA monoclonal antibody (mAb) (Sigma-Aldrich, H9658), rabbit anti-HA monoclonal antibody (mAb), mouse anti-Flag mAb (Sigma-Aldrich, F1804), mouse anti-Myc mAb (Sigma-Aldrich, M4439 and C3956), mouse anti-β-actin mAb (Sigma-Aldrich, A1978) were purchased from Sigma-Aldrich. Mouse anti-VP2 mAb of IBDV was produced and preserved in our laboratory. Anti-DFNA5/GSDME Rabbit Polyclonal Antibody (ER1901-12) was purchased from Hua Bio (Hangzhou, China). The mouse anti-Caspase-3 monoclonal antibody (mAb) (68773-1-Ig) was purchased from Proteintech (Wuhan, China).

### Enzyme-Linked Immunosorbent Assay

The level of IL-18 and IL-1β in cell culture supernatants were assayed using ELISA kits (SEA064Ga and SEA563Ga, Cloud-Clone). A total of 100 μL of standards or samples were added to corresponding wells, respectively. Then, 100 μL enzyme conjugate was added to the corresponding wells, sealed and incubated at 37 ℃ for 1 h. After washing, 50 μL of substrates A and 50 μL of B were added to each well and reacted at 37 ℃ for 15 min. Next, after adding 50 μL of stop solution, the optical density (OD) was read at 450 nm by a microtiter plate reader in 15 min. The cytokine concentrations could be calculated according to the standard curve.

### Co-immunoprecipitation (Co-IP) and western blotting

For co-immunoprecipitation (Co-IP), cells transfected with the indicated plasmids were lysed using NP-40 lysis buffer (Beyotime, P0013) and centrifuged at 12,000 × g for 10 minutes at 4℃. The whole cell lysates were pre-cleared with protein A/G agarose(Abmart, A10001) and subsequently incubated overnight at 4 ℃. The immunoprecipitated complexes were collected by centrifugation at 12,000 × g for 10 minutes at 4℃ and washed five times with PBS. After washing, the bound proteins were eluted by boiling in 5 x SDS loading buffer (Beyotime, P0015L) for 10 min and then subjected to western blotting analysis.

For western blotting analysis, the protein lysates were electrophoretically separated on 12.5% SDS-PAGE gel and then the proteins were transferred on nitrocellulose membranes. First, the membranes were blocked with 5% (w/v) skim milk in PBST at 37 ℃ for 1h, and then incubated with the corresponding primary antibody diluted with PBS. After washing four times with PBST, an appropriate incubated with an appropriate secondary antibody diluted with PBS was used. Finally, the membrane was washed four times in PBST and imaged using the Odyssey Infrared Imaging System (LICOR BioSciences, Lincoln, USA) for further analysis.

### Confocal microscopy

HEK-293T cells seeded in 35 mm dishes (Biosharp, BS-20-GJM) were co-transfected with the specified plasmids and incubated for 24h. Subsequently, the cells were rinsed three times with PBS and fixed using 4% (vol/vol) (Biosharp, BL539A) paraformaldehyde for 30 min. Following fixation, the cells were incubated with the appropriate primary antibodies, followed by incubation with the corresponding fluorescently labeled secondary antibodies. Finally, nuclei were counterstained with DAPI (Vectorlabs, H-1200) for 10 min, and images were acquired using a Leica SP2 confocal microscope (LSM980, Zeiss, Germany).

### RT-qPCR

Whole-cell RNA was extracted from transfected cells at indicated time points using the RNAiso Plus kit (9109, TaKaRa, Japan). The extracted RNA was reverse transcribed into cDNA using HiScript II QRT SuperMix for quantitative PCR (qPCR) (Vazyme, R223-01). The RT-qPCR amplification reaction utilized the SYBR Green qPCR Kit (TOYOBO, QPS-201) with the following cycling conditions host factors: initial denaturation at 95°C for 2 min, followed by 40 cycles of 95°C for 5 s, 60°C for 30 s, and followed by a melt curve analysis. The results were analyzed using the 2^-ΔΔCT^ method.

The primer sequences are as follows: *GSDMA* (F: 5’-AAGCACACCTTCA TCGAGCA-3’, R: 5’-GAGGAGATCACAGCGTGGAG-3’), *GSDME* (F: 5’-CAA AGGGCTGTGTGGAAAGC-3’, R: 5’-AGCCCCAGAGTACATCCCAT-3’), *β-acti n* (F: 5’-CAACACAGTGCTGTCTGGTGGTA-3’, R: 5’-ATCGTACTCCTGCTTGCTGATCC-3’), *Caspase-3* (F: 5’-CCATGGCGATGAAGGACTCT-3’, R: 5’-GTT TCCCTGCTAGACTTCTG-3’). *Caspase-9* (F: 5’-GGTCAAAGTTTGCCCTCTTTTCCTCATT-3’, R: 5’-GGAGCTGGGCCGCGCGGGCCCGGCCGGCCTGC-3’).

### Ethics statement

Specific pathogen-free (SPF) White Leghorn chickens were obtained from the Animal Experiment Center of HVRI, Chinese Academy of Agricultural Sciences. The animal experiments were approved by the Animal Ethics Committee of the HVRI (approval no. 240718-03-GR) and conducted in accordance with the international animal welfare standards (27).

### Animal experiment

Twenty-four 3-week-old specific-pathogen-free (SPF) chickens were randomly divided into two groups (12 per group), and intranasally inoculated with 1000 copies/200µL or an equal volume of PBS of IBDV SHG19, respectively. Clinical symptoms were monitored daily, and serum samples were collected at 1, 2, 3, 4, 5 and 6 days p.i, respectively. In addition, The bursa of two groups infected with SHG19 for 5 days p.i were fixed in 10% neutral buffered formalin, and histopathological examination was performed using hematoxylin and eosin (H&E) staining or scanning electron microscopy.

### Lactate dehydrogenase release assay

The LDH assay kit (Solarbio, BC0685) was purchased from Solarbio, the assay reagent was added according to the instructions from the manufacturer, and the optical density (OD) at 450 nm was read in the microplate reader.

### Drug treatment

In Caspase-3 activity inhibition experiments, Z-DEVD-FMK (Abmole, M3134) with a final concentration of 50ng/mL or dimethyl sulfoxide (DMSO) was added to the culture medium 2 h prior to SHG19 infection.

In Caspase-9 activity inhibition experiments, Z-LEHD-FMK (MBL, 4810-510) with a final concentration of 50ng/mL or DMSO was added to the culture medium 2h prior to SHG19 infection.

### Structure analysis

The structures of Caspase-9 and SHG19 VP3, SHG19S33A respectively were obtained from the Alphafold3 server, and the specific scoring structures were obtained. Further, the FastRelex module of Rosetta was used to optimize the protein-protein structure of the above three, obtaining the optimal structure of SHG19 VP3-Caspase-9. Meanwhile, Rosetta’s localing Protein-Protein docking module is used to further address the conflicts of interface amino acids and obtain a more reasonable protein complex structure. Finally, the InterfaceAnalysize module of Rosetta was used to further analyze the interfacial binding energy of each conformation in the three structures. Meanwhile, in the optimal structure of SHG19 VP3-Caspase-9, point mutation was used to convert it into the SHG19S33A protein, and Rosetta was used for evaluation. The interaction interface amino acids of the SHG19 VP3-Caspase-9 complex were analyzed by PLIP (28) and Ligplo (29).

### Statistical analysis

Data are presented as the mean ± standard deviation (SD). Statistical analyses were performed using GraphPad Prism software, including unpaired two-tailed Student’s *t* -tests, one-way ANOVA, or two-way ANOVA, followed by Dunnett’s multiple comparison test where appropriate. Error bars indicate the SD. Significance levels are denoted as follows: ns (not significant), *P* > 0.05; \**P* < 0.05; \*\**P* < 0.01; \*\*\**P* < 0.001.

## RESULTS

### nVarIBDV infection induces inflammatory responses associated with pyroptosis *in vivo* and *in vitro*

To evaluate the inflammatory properties induced by nVarIBDV infection, SPF chickens were infected with nVarIBDV (SHG19), and the bursae were collected for histopathological observation at 5 days post infection (dpi). The results of HE staining showed that the lymphoid follicles in the bursa were significantly atrophied, with a severe reduction in lymphocyte, and were accompanied by a large number of inflammatory cell infiltration (Fig 1A), indicating that acute inflammation is occurring in the bursa. To further detect the release of inflammatory factors, serum from chickens infected with SHG19 was collected for ELISA analysis. The results of ELISA showed that the expression levels of IL-1β (Fig 1B) and IL-18 (Fig 1C) were increased in the serum of the infected group from 1 dpi to 6 dpi compared to the mock group. Meanwhile, the results of RT-qPCR revealed that the transcription levels of IL-18 (Fig 1D) and IL-1β (Fig 1E) were also significantly upregulated in the SHG19 infected bursa. These results indicate that nVarIBDV infection induces severe inflammatory responses.

**FIG 1.**
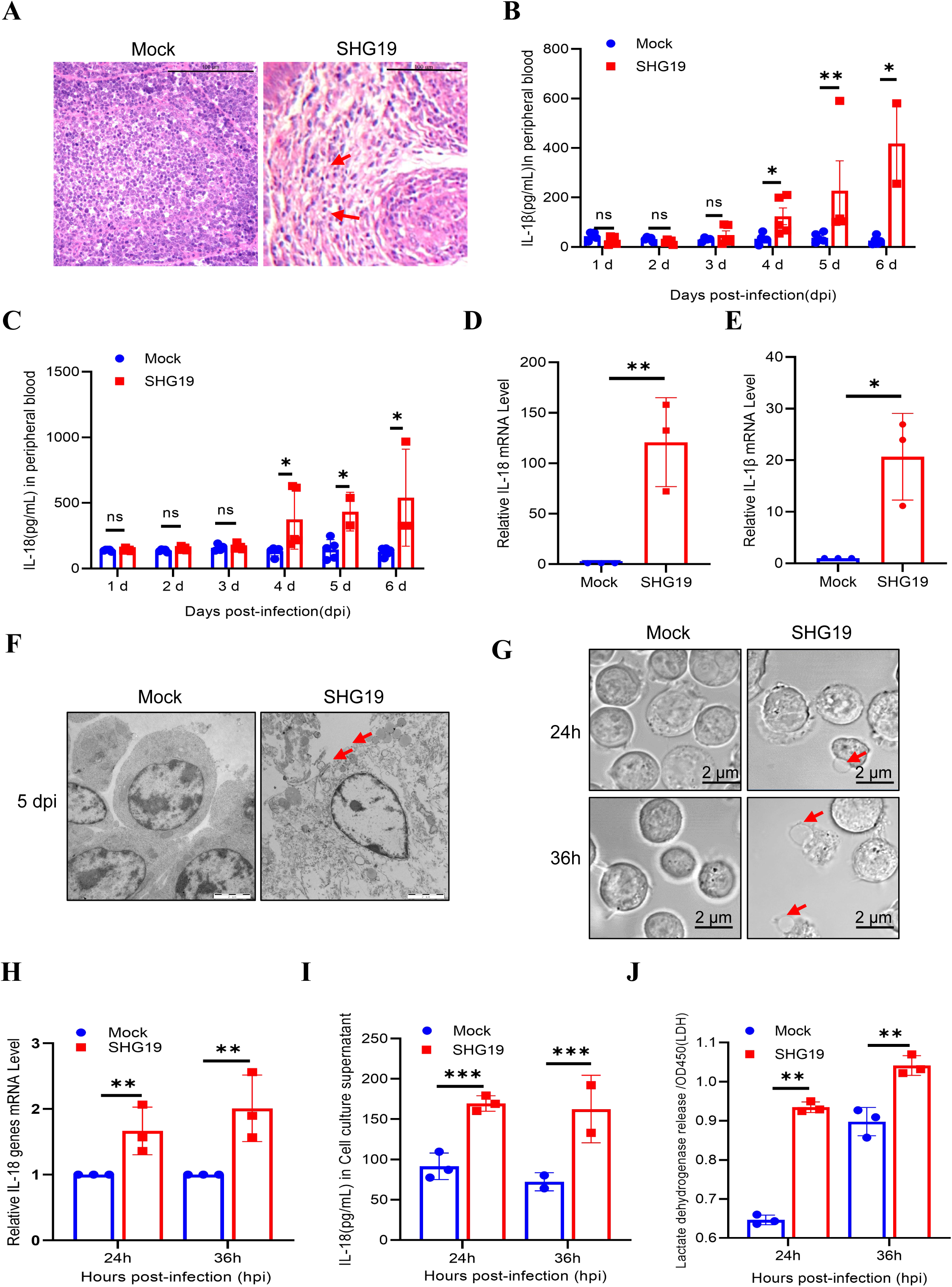
nVarIBDV infection induces inflammatory responses related to pyroptosis *in vivo* and *in vitro*. (A) Representative H&E staining analysis of the bursa sections of mock and nVarIBDV infected chickens at 5 dpi. The lymphoid follicles from nVarIBDV-infected chickens were severely atrophied with hemorrhage, lymphocyte depletion, and inflammatory cell infiltration. (B, C) Protein abundance of IL-18 (B) and IL-1β (C) in serums at indicated times post-infection was detected by ELISA. (D, E) Effects of SHG19 infection on IL-18 or IL-1β gene expression in the bursa. SPF chickens were infected with SHG19 (1×10^3^ copies/200 µl), and the mRNA levels of IL-18 (D) or IL-1β (E) were assessed by RT-qPCR at 5 dpi. (F) The structural changes of the bursa sections of mock and SHG19 infected chickens at 5 dpi were observed by Scanning electron microscope. Scale, 50 nm. Cell membrane fragmentation (red arrow). (G) Infection of DT40 cells with SHG19 induces pyroptosis. DT40 cells were infected with nVarIBDV at an MOI of 5. At 24 and 36 h post-infection (hpi), cells were observed under a confocal microscope, with arrows indicating pyroptotic cells. Scale, 2 µm. (H, I) Effects of SHG19 infection on IL-18 or IL-1β gene expression in DT40 cells. DT40 cells were infected with SHG19 (1×10^7^ copies/1×10^6^ cells), and the mRNA levels of IL-18 (H) or IL-1β (I) were assessed by RT-qPCR at 24 and 36 hpi. (J) Release of LDH from the infected cells was measured as described in Materials and Methods. Graphs show mean ± SD, n = 3, * *P* < 0.05, ** *P* < 0.01, *** *P* < 0.001.

Pyroptosis serves as both an “executor” and “amplifier” of inflammation, characterized by cell swelling and membrane bubbling, which leads to the passive release of IL-1β and IL-18. To further verify whether the inflammatory responses is related to pyroptosis, SHG19-infected bursae were fixed and examined by scanning electron microscopy. These results showed that SHG19 infection could cause blurred cellular morphology and disrupted cell membranes within the bursa, similar to the characteristics of pyroptosis (Fig 1F). To further confirm the above results *in vitro*, DT40 cells were infected with SHG19 at a multiplicity of infection (MOI) of 5, the results of confocal microscopy showed that DT40 cells exhibited typical features of pyroptosis (swelling and rupture of cell membrane) at 24- and 36- hours post infection (hpi) (Fig 1G). In addition, RT- qPCR results showed that DT40 cells infected with SHG19 were also accompanied by a significant increase in the transcription levels of IL-18 (Fig 1H). Meanwhile, the results of ELISA similarly showed that the protein level of IL-18 in the supernatant was significantly increased (Fig 1I) and LDH release was also significantly increased after SHG19 infection (Fig 1J). In conclusion, these results suggest that inflammation induced by nVarIBDV is associated with pyroptosis.

### nVarIBDV induces pyroptosis through the Caspase-3-GSDME pathway

The implementation of pyroptosis depends on the oligomerization of the N-terminal domain of GSDM family proteins on the cell membrane after they are cleaved by proteases. The members of this family in chickens are quite special, the GSDMD gene is missing, and currently only GSDMA and GSDME are known to be active full-length proteins (30, 31). To determine the possible pathway by which nVarIBDV induces pyroptosis, we detected the expression levels of pyroptosis-related proteins after SHG19 infection *in vivo* and *in vitro*, respectively. The results of RT-qPCR showed that the transcription level of GSDME was significantly upregulated, whereas no significant change was observed in the transcription level of GSDMA in the bursa infected with SHG19 (Fig 2A). Similarly, a significant increase in the transcription level of GSDME was also observed in DT40 cells at 24 hpi, compared with GSDMA (Fig 2B). Meanwhile, the results of western blotting also indicated that GSDME was significantly cleaved in DT40 cells after SHG19 infection (Fig 2C). These results indicate that SHG19 infection induced pyroptosis mainly through GSDME. Caspase-3 specifically cleaves GSDME to release its pore-forming N-terminal domain, which triggers the execution of pyroptosis (32). To verify whether SHG19 infection induces pyroptosis through the Caspase-3-GSDME pathway, we examined the activation of Caspase-3 after SHG19 infection. The results of western blotting revealed that Caspase-3 was significantly activated after SHG19 infection (Fig 2C), suggesting that SHG19 infection-induced pyroptosis might be executed through the Caspase-3-GSDME pathway. To further confirm that Caspase-3 is essential for GSDME-mediated pyroptosis following SHG19 infection, DT40 cells were infected with SHG19 either in the presence or absence of Z-DEVD-FMK (a Caspase-3-specific inhibitor), and pyroptosis-related indicators were subsequently detected. As shown in Fig 2D, the results of confocal microscopy showed that DT40 cells treated with Z-DEVD-FMK did not exhibit typical pyroptosis features. The western blotting results showed that the GSDME-N fragment induced by SHG19 infection was significantly reduced in the presence of Z-DEVD-FMK (Fig 2E). Meanwhile, the release of IL-18 (Fig 2F) and LDH (Fig 2G) induced by SHG19 infection were also significantly inhibited in the presence of Z-DEVD-FMK. These results demonstrated that SHG19 infection induce pyroptosis and inflammation through the Caspase-3-GSDME pathway.

**FIG 2.**
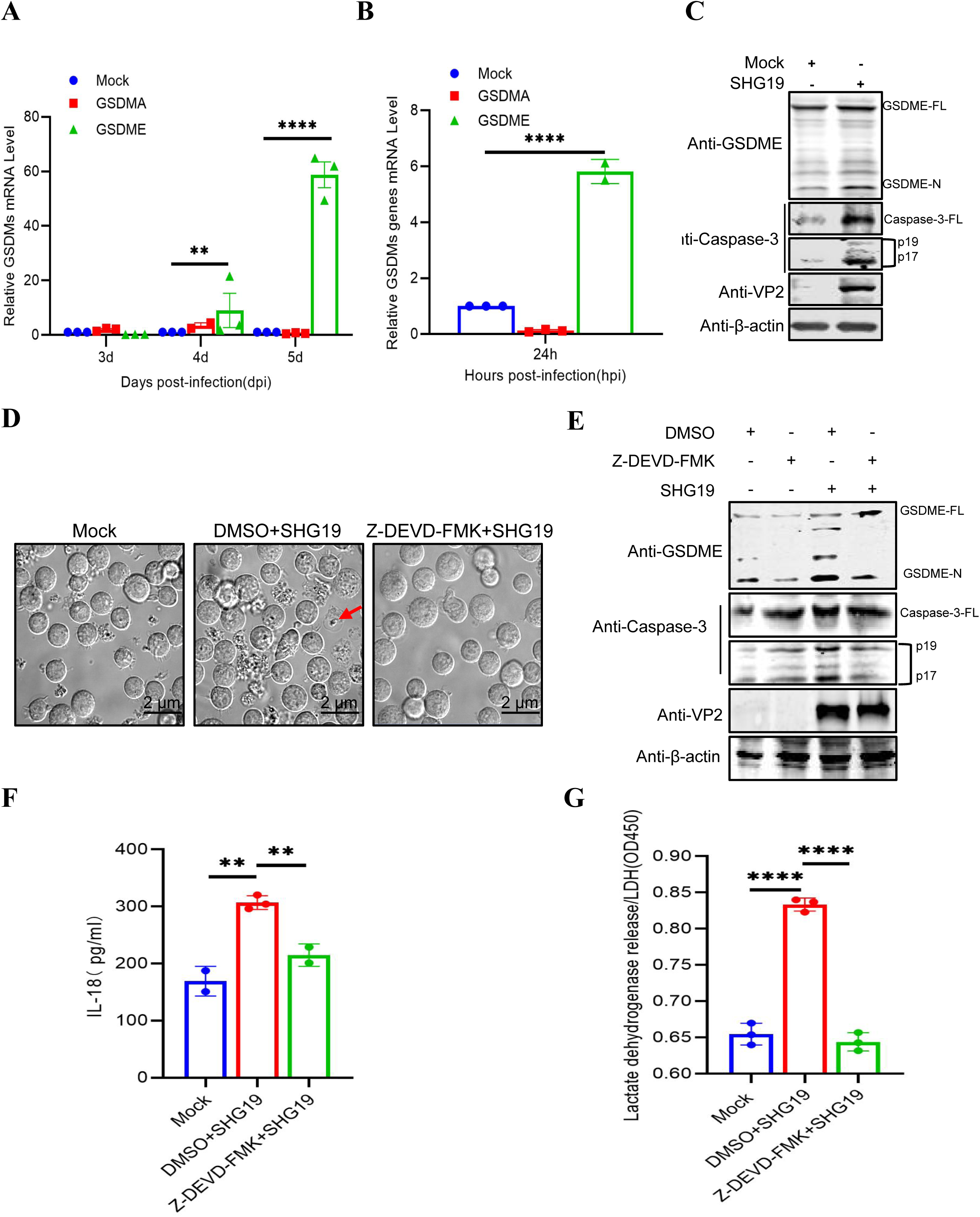
nVarIBDV infection induces pyroptosis in DT40 cells dependent on Caspase-3 activity. (A) Effects of SHG19 infection on GSDMA or GSDME gene expression in the bursa. SPF chickens were infected with SHG19 (1×10^3^ copies/200 µl), and the mRNA levels of GSDMA or GSDME were assessed by RT-qPCR at 3, 4 and 5 dpi. (B) Effects of SHG19 infection on GSDMA or GSDME gene expression in DT40 cells. DT40 cells were infected with SHG19 (1×10^7^ copies/1×10^6^ cells), and the mRNA levels of GSDMA or GSDME were assessed by RT-qPCR at 24 and 36 hpi. (C) SHG19 infection induces GSDME-mediated pyroptosis in DT40 cells. Proteolytic cleavage of GSDME in SHG19-infected DT40 cells was determined by western blotting at 24 hpi. The abundance of Caspase-3, SHG19 VP2 protein, and β-actin as an internal control was also determined by western blotting. (D) Confocal microscopy observation of pyroptosis. DT40 cells were infected with SHG19 at an MOI of 5 in the presence or absence of the Z-DEVD-FMK, DT40 cells were observed under a confocal microscope, with arrows indicating pyroptotic cells. Scale, 2 µm. (E) Proteolytic cleavage of GSDME in DT40 cells with SHG19 infection was determined by western blotting at 24 hpi in the presence or absence of the Z-DEVD-FMK (Caspase-3-specific inhibitor). (F, G) Protein abundance of IL-18 (F) and LDH (G) in cell culture supernatant at indicated times post-infection in the presence or absence of the Z-DEVD-FMK were detected by ELISA. Graphs show mean ± SD, n = 3, * *P* < 0.05, ** *P* < 0.01, **** *P* < 0.0001.

### Viral protein VP3 of nVarIBDV activates Caspase-3 and mediates GSDME cleavage to induce pyroptosis

To screen the key viral protein responsible for SHG19-induced pyroptosis, the pCAGGS-HA eukaryotic expression plasmids of SHG19 viral proteins (VP1, VP2, VP3, VP4 and VP5, respectively) were co-transfected with the pMyc-Caspase-3 and pFlag-GSDME (with Flag tags fused at both N- and C- terminal) into HEK-293T cells, the cleavage of GSDME and LDH release were detected. As shown in Fig 3A, viral protein VP3 enhanced the cleavage of GSDME of Caspase-3-mediated, while other viral proteins did not. Consistently, LDH assays verified that VP3 significantly increased the release of LDH mediated through the Caspase-3-GSDME pathway (Fig 3B). To verify the above results, we transfected HEK-293T cells with pMyc-Caspase-3, pFlag-GSDME and different doses of pHA-VP3 (0, 1, 2μg, respectively) to detect the cleaved N-terminal fragment of GSDME. The results of western blotting showed that VP3 promoted the cleavage of GSDME of Caspase-3-mediated in a dose-dependent manner (Fig 3C).

**FIG 3.**
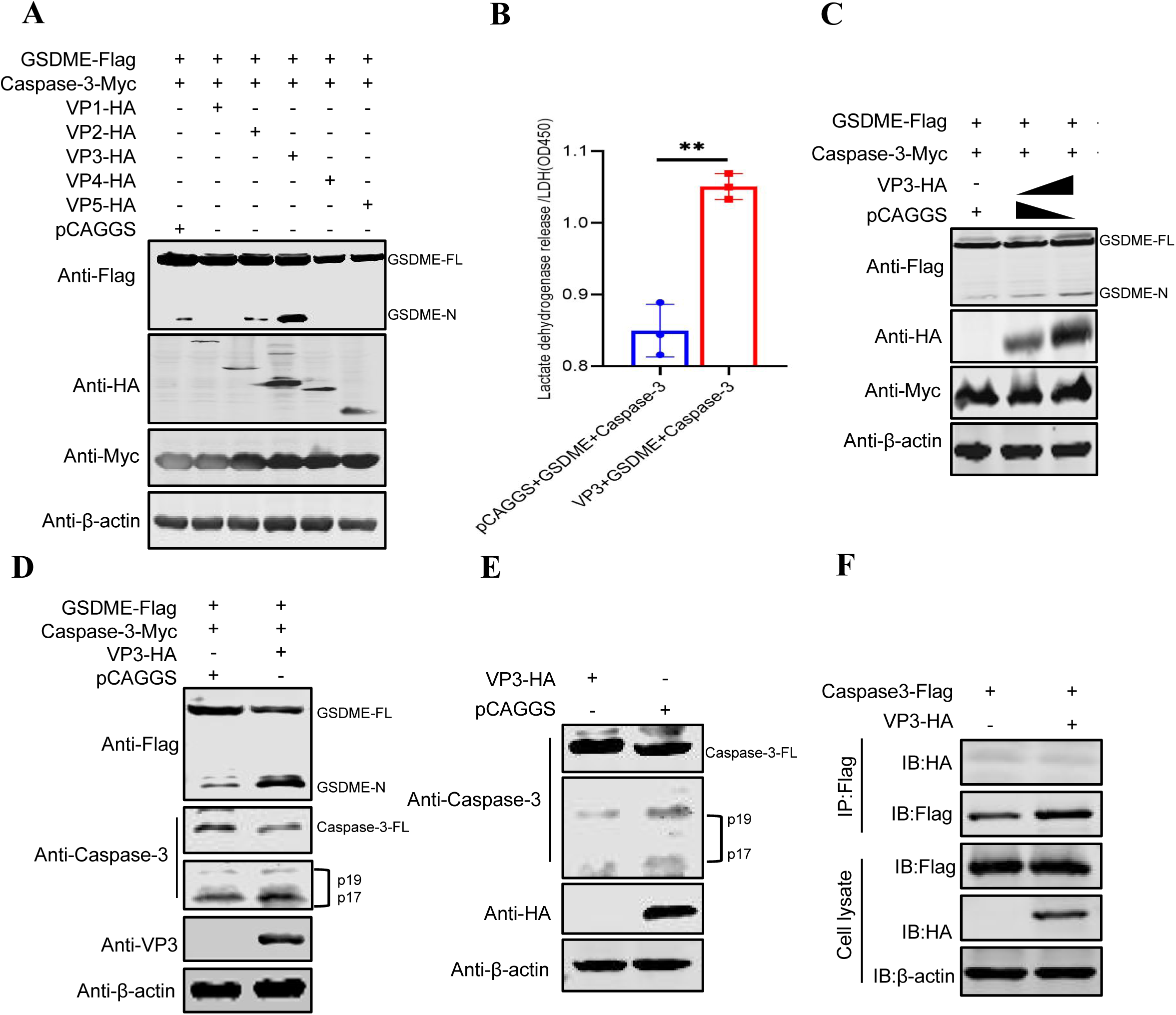
viral protein VP3 induces pyroptosis by promoting Caspase-3 activation to cleave GSDME. (A) VP3 promotes Caspase-3-mediated cleavage of GSDME. HEK-293T cells were co-transfected with pMyc-Caspase-3, pFlag-GSDME and eukaryotic plasmids expressing viral protein-HA fusions containing VP1, VP2, VP3, VP4, or VP5 of SHG19. Twenty-four hours post-transfection, cell lysates were examined by western blotting using anti-Myc, anti-HA, anti-Flag, and anti-β-actin antibodies; endogenous β-actin expression was used as an internal control. (B) LDH release from the cells transfected with pMyc-Caspase-3, pFlag-GSDME and pHA-VP3 was measured as described in Materials and Methods. (C) VP3 promotes Caspase-3 cleavage of GSDME in a dose-dependent manner. HEK-293T cells were co-transfected with pMyc-Caspase-3, pFlag-GSDME and different concentrations of pHA-VP3 to detect the cleavage of GSDME using western blotting. (D, E) VP3 activates both endogenous and exogenous Caspase-3. HEK-293T cells were co-transfected with pMyc-Caspase-3, pFlag-GSDME and pHA-VP3, or VP3 alone. Twenty-four hours post-transfection, cell lysates were examined with western blotting anti-Myc, anti-HA, anti-Flag, anti-Caspase-3 and anti-β-actin antibodies. (F) The interaction between VP3 and Caspase-3 detected by Co-IP. HEK-293T cells were transfected with indicated plasmid and harvested at 30 hpi. Cell lysates were immunoprecipitated using Flag Tag and analysed using the HA Tag and Flag Tag. Graphs show mean ± SD, n = 3, ** *P* < 0.01.

Next, we investigated the mechanism by which VP3 enhances the cleavage of GSDME of Caspase-3-mediated by assessing its influence on both exogenous and endogenous Caspase-3 activation. The results of western blotting showed that VP3 significantly promoted the activation of exogenous Caspase-3 during its mediation of GSDME cleavage (Fig 3D). Consistently, VP3 also potently activated endogenous Caspase-3 (Fig 3E). To further verify whether VP3 promotes Caspase-3 activation through direct interaction with Caspase-3, HEK-293T cells were co-transfected with plasmids expressing pFlag-Caspase-3 and either pHA-VP3 or pCAGGS. The results of co-immunoprecipitation (Co-IP) indicated that VP3 did not interact with Caspase-3 directly (Fig 3F). In conclusion, these results indicate that VP3 indirectly promotes the activation of Caspase-3 and further cleavage of GSDME, thereby inducin<u>g</u> pyroptosis.

### VP3 interacts with Caspase-9 to activate Caspase-3-GSDME pathway to induce pyroptosis

Previous studies have shown that Caspase-3 can be activated by the upstream molecule Caspase-9, a core protein of the endogenous apoptotic pathway (33). To verify whether VP3 promotes the activation of Caspase-3 through Caspase-9, the expression level of Caspase-9 was detected after SHG19 infection or transfection of VP3 into DF-1 cells, respectively. The results of RT-qPCR showed that the transcription levels of Caspase-9 were significantly increased after SHG19 infection at 24 and 36 hpi, respectively (Fig 4A). Meanwhile, transfection of DF-1 cells with pHA-VP3 also increased the transcription level of Caspase-9 (Fig 4B). These results suggest that Caspase-9 may be a key target regulated by VP3.

**FIG 4.**
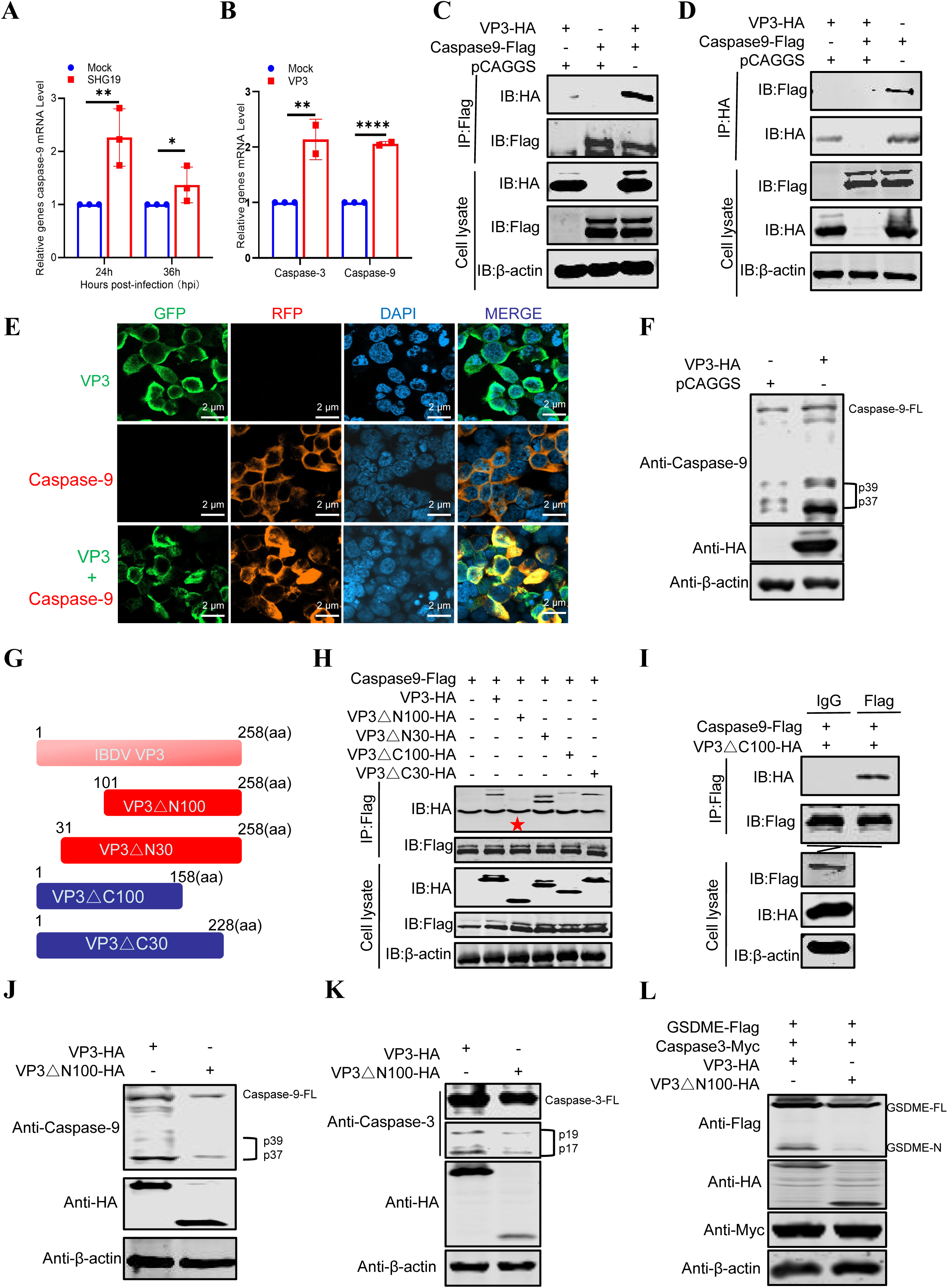
VP3 interacts with Caspase-9 to promote activation of Caspase-9. (A) Effects of nVarIBDV infection on Caspase-3 and Caspase-9 gene expression in DT40 cells. DT40 cells were infected with SHG19 (1×10^7^ copies/1×10^6^ cells), and the mRNA levels of Caspase-3 and Caspase-9 were assessed by RT-qPCR at 24 and 36 hpi. (B) Effects of VP3 transfection on Caspase-3 and Caspase-9 gene expression in DF-1 cells. After DF-1 cells were transfected with VP3 or pCAGGS for 24h, the cells were collected to detect the mRNA levels of Caspase-3 and Caspase-9. All data are representative of at least three independent experiments. Graphs show mean ± SD, n = 3, * *P* < 0.05, ** *P* < 0.01, *** *P* < 0.001. (C, D) The interaction between VP3 and Caspase-9 as detected by Co-IP. (E) Confocal assays were used to assess the colocalization between Caspase-9 and VP3. HEK-293T cells were co-transfected with pFlag-Caspase-9 and pHA-VP3 for 36h. Cells were incubated with anti-Flag mAb produced in mice and anti-HA mAb produced in rabbits and the interaction between Caspase-9 and VP3 was determined by confocal analysis. (F) VP3 promotes the activation of Caspase-9. DF-1 cells were transfected with pCAGGS or VP3. Twenty-four hours post-transfection, cell lysates were examined with western blotting using anti-HA, anti-Caspase-9, and anti-β-actin antibodies; endogenous β-actin expression was used as an internal control. (G) Schematic representation of the structure of VP3 and its mutants. (H, I) The interaction between VP3, VP3ΔN100, VP3ΔC100, VP3ΔN30, VP3ΔC30 and Caspase-3 as detected by Co-IP. (J-K) The influence of VP3 and VP3ΔN100 on the activation of Caspase-9 and Caspase-3. DF-1 or HEK-293T cells were transfected with pHA-VP3ΔN100 or pHA-VP3. Twenty-four hours post-transfection, cell lysates were examined with western blotting using anti-HA, anti-Caspase-9, anti-Caspase-3 and anti-β-actin antibodies. (L) Assessment of the ability of VP3ΔN100 to promote Caspase-3 to cleave GSDME. HEK-293T cells were co-transfected with pMyc-Caspase-3, pFlag-GSDME and VP3 or VP3ΔN100, cell lysates were examined with western blotting using anti-Myc, anti-HA, anti-Flag, and anti-β-actin antibodies. Graphs show mean ± SD, n = 3, ** *P* < 0.01, **** *P* < 0.0001.

To determine whether VP3 interacts with Caspase-9, we co-transfected pFlag-Caspase-9 and pHA-VP3 into HEK-293T cells to detect their interaction. The results of Co-IP indicated that VP3 strongly interacted with Caspase-9 (Fig 4C). Similarly, as shown in Fig 4D, the reverse Co-IP results verified that Caspase-9 indeed interacted with VP3. Confocal microscopy results indicated that Caspase-9 co-localized with VP3 in the cytoplasm (Fig 4E). Caspase-9 needs to undergo auto-cleavage to transform into the active forms to activate Caspase-3. To determine whether VP3 can promote the activation of Caspase-9, pHA-VP3 was transfected into DF-1 cells, and the active forms of Caspase-9 were examined. As shown in Fig 4F, VP3 significantly promoted the activation of Caspase-9, converting full-length Caspase-9 to p37 or p39 fragments.

To further confirm that the interaction between VP3 and Caspase-9 is essential for activating the Caspase-9-Caspase-3-GSDME-mediated pathway, we disrupted this interaction by generating a binding-deficient VP3 mutant and assessed the subsequent activation of Caspase-9, Caspase-3, and the cleavage of GSDME. First, four VP3 truncated mutants (pHA-VP3 Δ N30, pHA-VP3 Δ N100, pHA-VP3 Δ C100 and pHA-VP3ΔC30) were constructed (Fig 4G), and the ability of each to interact with Caspase-9 was examined. The results of Co-IP showed that VP3ΔC30, VP3ΔC100, and VP3ΔN30 mutants interacted with Caspase-9, whereas VP3ΔN100 showed no interaction (Fig 4H and 4I). These results indicated that the interaction between VP3 and Caspase-9 was located within the N-terminal 30-100 amino acid (aa) region. To determine whether the N-terminal of VP3 was crucial for promoting the activation of Caspase-9, we transfected pHA-VP3 or pHA-VP3ΔN100 into DF-1 cells to detect their ability to activate Caspase-9. The results of western blotting showed that the deletion of 30-100 aa at the N-terminal of VP3 significantly reduced the ability to activate Caspase-9 (Fig 4J). Correspondingly, VP3ΔN100 significantly weakened the ability to activate Caspase-3 (Fig 4K) and to cleave GSDME (Fig 4L) compared to the wild type (WT) VP3. In summary, VP3 activates Caspase-9 by interacting with it, thereby promoting the cleavage of GSDME mediated by Caspase-3 to induce pyroptosis.

### Ser33 of VP3 is the key amino acid that activates Caspase-9 to induce pyroptosis

To further identify the key amino acid of VP3 involved in interacting with Caspase-9, another three truncated mutants of VP3 (pHA-VP3Δ40, pHA-VP3Δ60 and pHA-VP3Δ80) located in the N-terminal 30-100 aa region were constructed. The results of Co-IP showed that all three truncates of VP3 lost the ability to interact with Caspase-9 (Fig 5A), indicating that the key region interacting with Caspase-9 is located within the 31-39 aa region at the N-terminal. To further narrow down the range, a mutant lacking the N-terminal 35 aa (pHA-VP3Δ35) was constructed.

**FIG 5.**
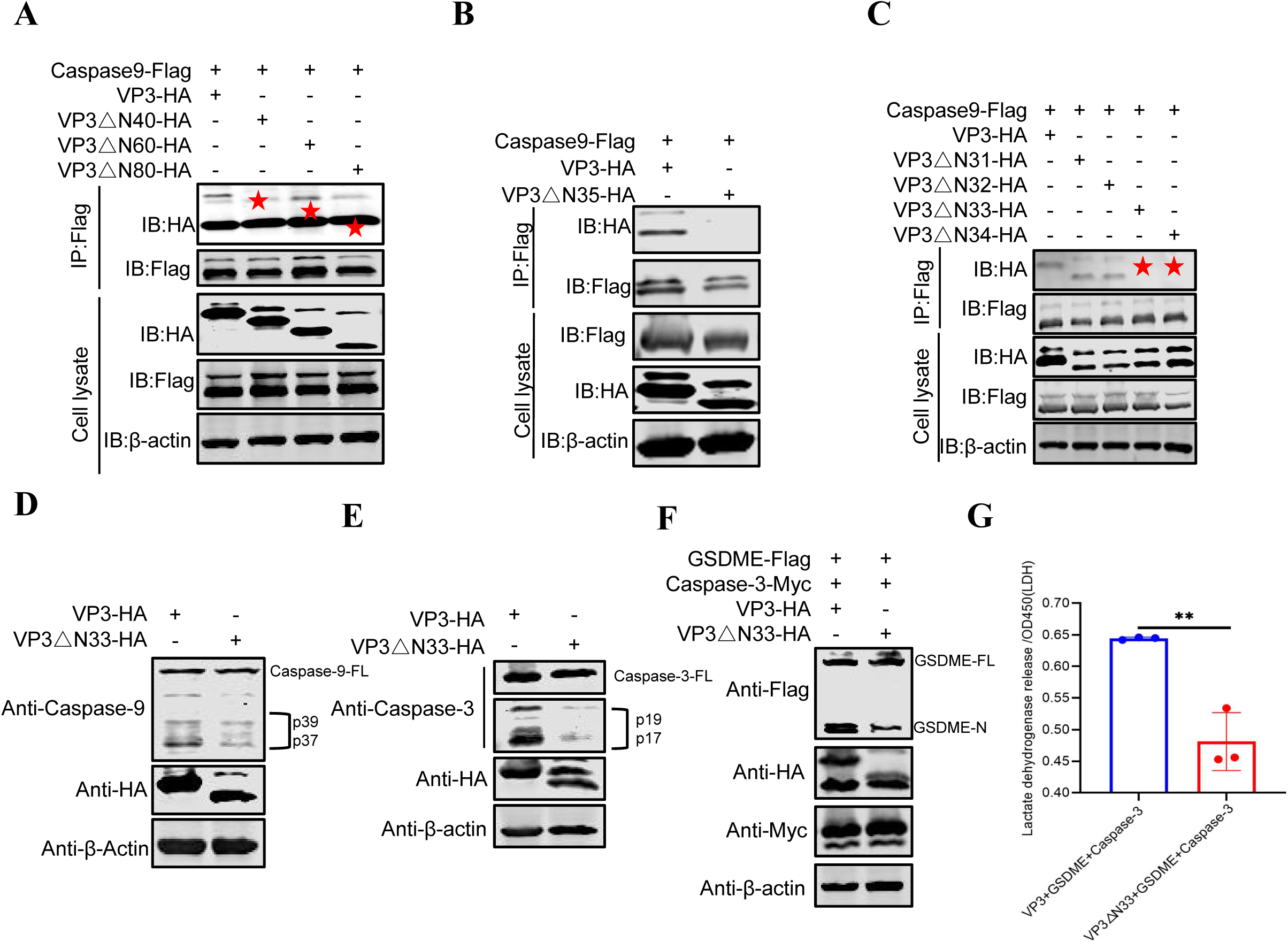
Exploring the key amino acid sites mediating the interaction between VP3 and Caspase-9. (A-C) The interaction between VP3, VP3Δ40, VP3ΔN60, VP3 ΔN80, VP3ΔN35, VP3ΔN31, VP3ΔN32, VP3ΔN33, VP3ΔN34 and Caspase-9 as detected by Co-IP as described above. (D) Effects of of VP3 and VP3ΔN33 on the activation of the Caspase-9 and Caspase-3. DF-1 or HEK-293T cells were transfected with VP3ΔN100 or VP3. Twenty-four hours post-transfection, cell lysates were examined by western blotting using anti-HA, anti-Caspase-9, anti-Caspase-3 and anti-β-actin antibodies. (F) Assessment of the ability of VP3ΔN33 to promote Caspase-3 to cleave GSDME. (G) Release of LDH from the cells transfected with pMyc-Caspase-3, pFlag-GSDME and pHA-VP3 or pHA-VP3ΔN33 was measured as described in Materials and Methods section. Graphs show mean ± SD, n = 3, ** *P* < 0.01.

Further analysis confirmed that the absence of 35 aa of VP3 did not interact with Caspase-9 (Fig 5B), indicating the key region limited as 31-34 aa. Therefore, four VP3 truncated mutants (pHA-VP3 Δ 31, pHA-VP3 Δ 32, pHA-VP3 Δ 33 and pHA-VP3 Δ 34) were constructed and co-transfected with pFlag-Caspase-9 into HEK-293T cells. The results of Co-IP showed that both the pHA-VP3Δ33 and pHA-VP3Δ34 mutants failed to interact with pFlag-Caspase-9, but pHA-VP3Δ31 and pHA-VP3Δ32 mutants interacted with pFlag-Caspase-9, indicating that VP3 residue 33 (Ser33) constituted a critical site for interacting with Caspase-9 (Fig 5C). To further validate the importance of the Ser33 of VP3 in activating Caspase-9, HEK-293T cells were transfected with either the pHA-VP3Δ33 truncated mutant or pHA-VP3 to detect their ability to activate Caspase-9 and Caspase-3. The results of western blotting indicated that VP3Δ33 truncated mutant significantly reduced the activation of Caspase-9 (Fig 5D) and Caspase-3 (Fig 5E). Correspondingly, its ability to promote Caspase-3-mediated cleavage of GSDME was also significantly impaired (Fig 5F). Compared with WT VP3, the release of LDH was also significantly reduced after deletion of residue 33 in VP3 (Fig 5G). These results indicate that residue 33 of VP3 (Ser33) plays a significant role in the Caspase-9-Caspase-3-GSDME pathway.

### Mutation at Ser33 of VP3 weakens the structural basis of Caspase-9 binding ability

To further verify the significance of Ser33 in its interaction with Caspase-9, we constructed a mutant plasmid in which serine 33 of VP3 was mutated to Alanine (Ala) (pHA-VP3S33A), and examined its ability to bind to Caspase-9 and activate Caspase-9 and Caspase-3. The results showed that, compared with WT VP3, pHA-VP3S33A not only exhibited a significantly weakened interaction with pFlag-Caspase-9 in Co-IP assays (Fig 6A) but also a markedly reduced ability to activate endogenous Caspase-9 (Fig 6B) and Caspase-3 (Fig 6C). The active forms of Caspase-9 and Caspase-3 were significantly reduced. To further verify the effect of pHA-VP3S33A on VP3-induced pyroptosis, pHA-VP3S33A or pHA-VP3 were co-transfected with the pMyc-Caspase-3 and pFlag-GSDME into the HEK-293T cells and the cleavage of GSDME and LDH release were detected. As expected, the ability of pHA-VP3S33A to promote cleavage of GSDME of Caspase-3-mediated was significantly reduced compared with pHA-VP3 (Fig 6D), ultimately leading to a corresponding decrease in LDH release (Fig 6E). Collectively, our results demonstrated that residue 33 of VP3 is a key amino acid for Caspase-9 activation and played a crucial role in activating the Caspase-9-Caspase-3-GSDME pathway and ultimately leading to GSDME cleavage.

**FIG 6.**
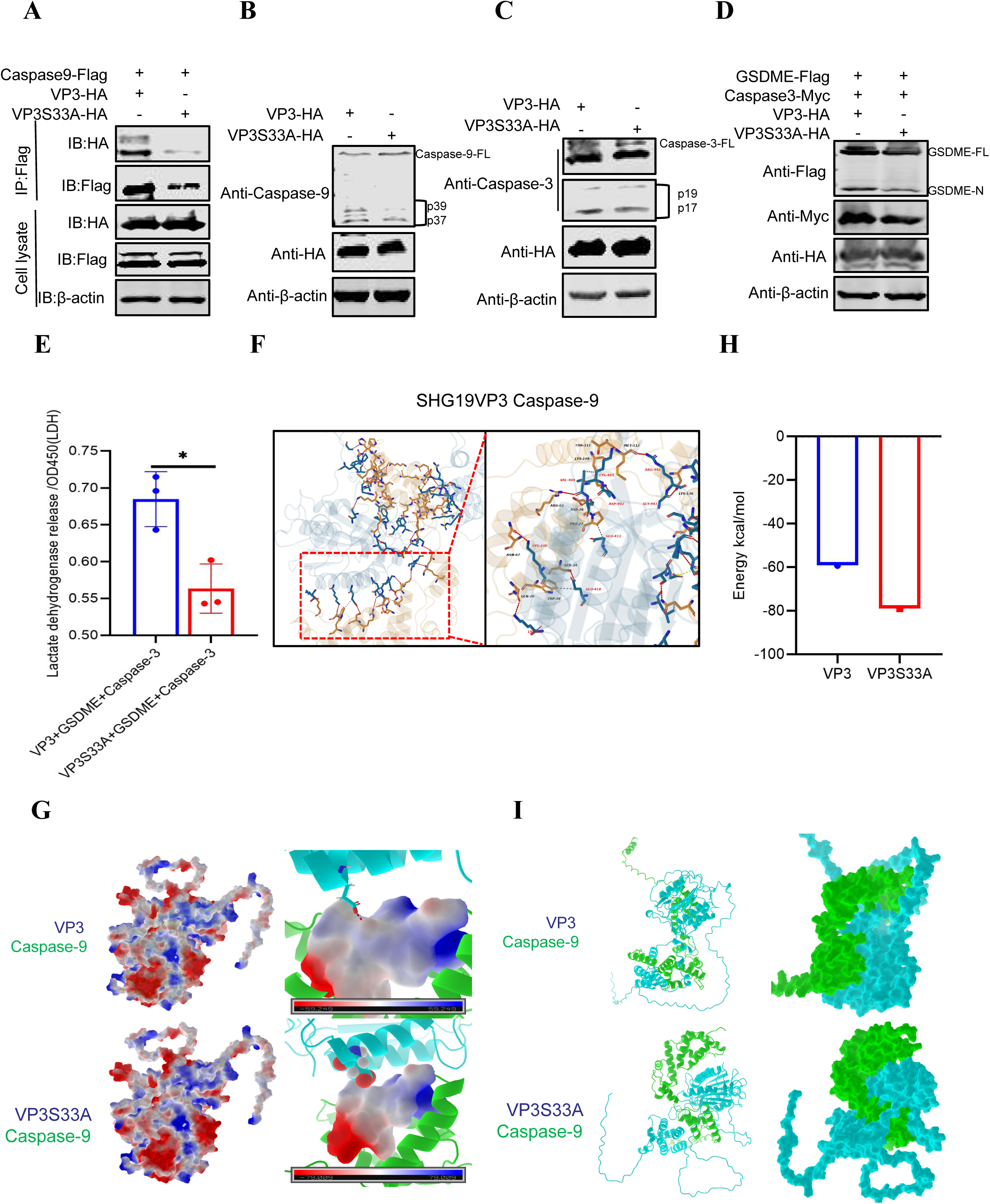
Effect of the Ser33 mutation on VP3 activation of Caspase-9-Caspase-3-GSDME pathway. (A) The interaction between VP3, VP3S33A and Caspase-9 as detected by Co-IP. (B, C) Effects of VP3 and VP3S33A on the activation of the Caspase-9 and Caspase-3 as determined by western blotting. (D) Assessment of the ability of VP3S33A to promote Caspase-3 to cleave GSDME. (E) LDH release induced by VP3 and VP3S33A via the Caspase-3-GSDME pathway. (F) Structural model of the interaction between VP3 and Caspase-9. The structure and surface of SHG19 VP3-Caspase-9 (SHG19 VP3 Chain A shown in wheat, Caspase-9 Chain B show in slate). The SHG19 VP3-Caspase-9 interactions (Caspase-9 key residues shown in red, Hydrogen Bonds shown in red, Hydrophobic Interactions shown in white, Salt Bridges shown in yellow). (G) The changes in the surface electrostatic potential of the binding motif after VP3 and VP3S33A bind to Caspase-9. The electrostatic potential was calculated and visualized with blue indicating positive potential and red indicating negative. (H) Comparison of the binding free energies of VP3 and VP3S33A binding to Caspase-9. (I) The structure comparison of VP3 and VP3S33A in complex with Caspase-9 (VP3 and VP3S33A shown in green, and Caspase-9 shown in cyan). Graphs show mean ± SD, n = 3, * *P* < 0.05.

To gain mechanistic insight into how Ser33 mediates the VP3-Caspase-9 interaction identified above, we performed molecular docking to simulate their binding mode. These results showed that the binding free energy at the interface between VP3 and Caspase-9 was -124.243 kcal/mol, and Ser 33 of VP3 formed a hydrogen bond with Glu418 of Caspase-9 (Fig 6F), indicating that the interaction between VP3 and Caspase-9 was highly reliable. To further assess the effect of the Ser33 mutation on VP3-Caspase-9 binding, we used AlphaFold3 to model the complexes and analyze their interfacial electrostatic potentials, visualized as alternating positive (blue) and negative (red) patches. The results indicated that mutation at Ser33 of VP3 made the disappearance of the hydrogen bond at this position of bingding with Caspase-9 disappeared (Fig 6G), and also changed the electrostatic potential of this binding site from -59.249 kcal/mol to -79.009 kcal/mol (Fig 6H), thereby increasing the instability of the binding of VP3 protein to Caspase-9. It is worth noting that the interaction structure between VP3S33A and Caspase9 was significantly different from that between VP3 and Caspase-9 (Fig 6I), indicating that the Ser33 of VP3 plays a significant role in maintaining the stable conformation of Caspase-9 in the VP3-Caspase-9 protein-interaction mode.

### nVarIBDV induces pyroptosis and inflammatory responses by directly activating Caspase-9

To further determine the key role of Caspase-9 in the process of pyroptosis and inflammation induced by nVarIBDV, we infected DT40 cells with SHG19 at a MOI of 5 for 12 hours to detect pyroptosis-related indicators in the presence or absence of the Z-LEHD-FMK (Caspase-9-specific inhibitor). The results of western blotting showed that in the presence of Z-LEHD-FMK, the ability of SHG19 infection to activate endogenous Caspase-3 was significantly inhibited, and accordingly, the cleavage fragment of GSDME was significantly reduced (Fig 7A). Z-LEHD-FMK significantly inhibited endogenous Caspase-9 expression (Fig 7B). Further, the results of confocal microscopy revealed that DT40 cells treated with Z-DEVD-FMK did not exhibit typical features of pyroptosis (Fig 7C). Moreover, the LDH release induced by SHG19 infection was significantly reduced in the presence of Z-LEHD-FMK (Fig 7D). As expected, the release of IL-18 after SHG19 infection was also significantly reduced compared with the mock group in the presence of Z-LEHD-FMK (Fig 7E). These results demonstrate that the nVarIBDV-induced pyroptosis and inflammatory responses are dependent on the activation of Caspase-9.

**FIG. 7.**
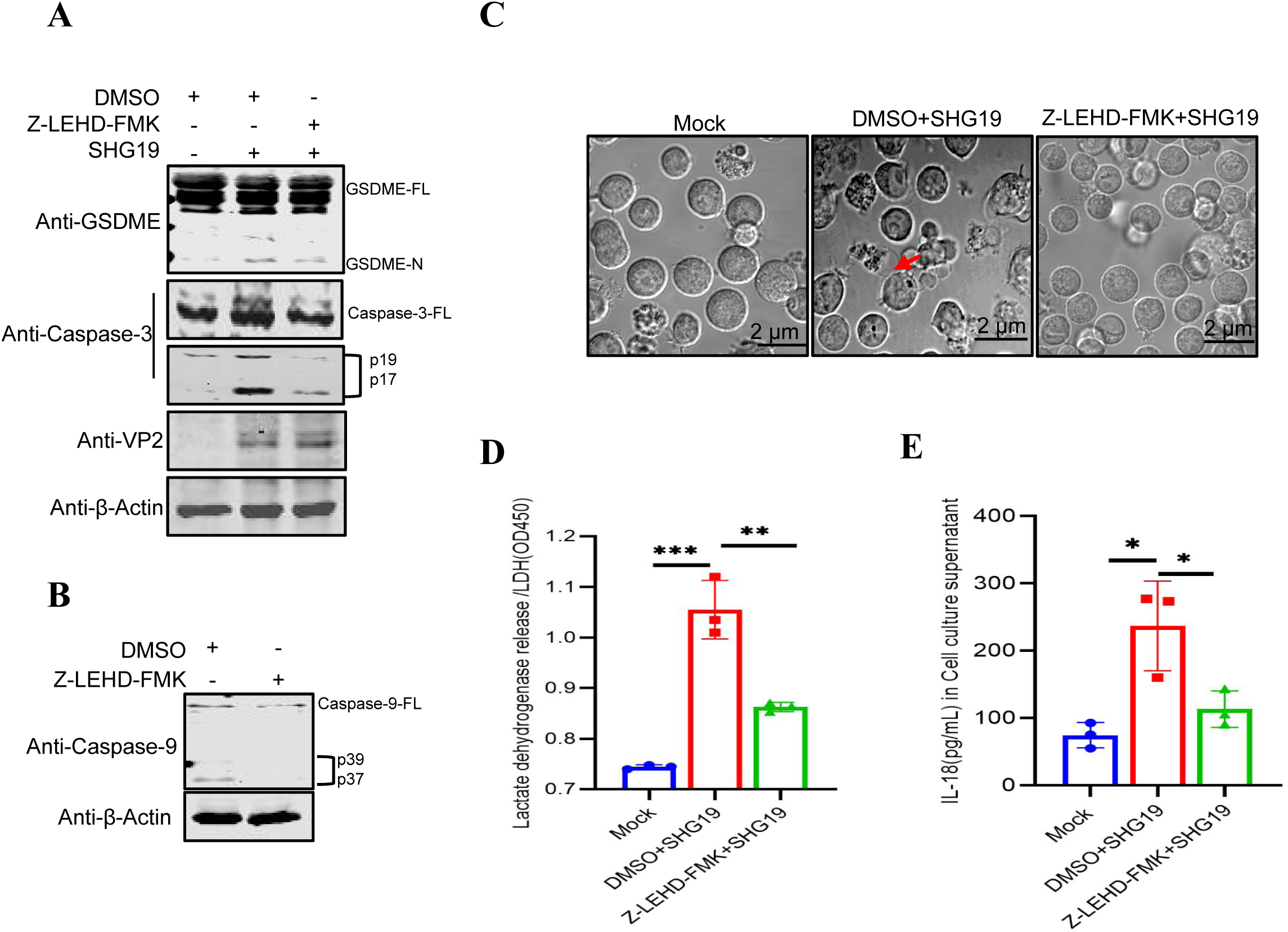
The effect of Caspase-9 inhibitors on pyroptosis of B lymphocyte induced by nVarIBDV. (A) Proteolytic cleavage of GSDME in DT40 cells following SHG19 infection was determined by western blotting at 24 hpi in the presence or absence of the Z-LEHD-FMK. (B) Detection of the inhibitory effect of Z-LEHD-FMK on endogenous Caspase-9 in DF-1 cells. (C) Confocal microscopy analysis of pyroptosis. DT40 cells were infected with SHG19 at an MOI of 5 in the presence or absence of the Z-LEHD-FMK, and examined by confocal microscopy, with arrows indicating pyroptotic cells. (D, E) Protein abundance of LDH (D) and IL-18 (E) in the cell supernatant at 24 hpi in the presence or absence of the Z-LEHD-FMK was measured by ELISA. Graphs show mean ± SD, n = 3, * *P* < 0.05, ** *P* < 0.01, *** *P* < 0.001.

## DISCUSSION

Since nVarIBDV emerged, it has become predominant circulating strain worldwide, posing a serious threat to the poultry industry (34, 35). Although it differs from the previously prevalent vvIBDV in terms of virulence and clinical pathogenic features, they share a common core pathogenic mechanism: triggering severe inflammatory responses and causing irreversible damage to the bursa of Fabricius (14). However, the molecular mechanisms underlying nVarIBDV-induced inflammatory pathogenesis remain largely elusive, which significantly hampers the development of effective prevention and control strategies. This study demonstrates that nVarIBDV induces inflammation through a novel pathway with VP3-Caspase-9 as the core axis, whereby VP3 directly activates Caspase-9, thereby driving Caspase-3-GSDME-mediated pyroptosis of B lymphocyte (Fig 8). Further mechanism studies have shown that the Ser33 residue of VP3 is the key site for its binding to Caspase-9 and for maintaining Caspase-9 active conformation. This discovery reveals the molecular basis by which nVarIBDV precisely regulates the host cell death pathway to induce pyroptosis and inflammation.

**FIG. 8.**
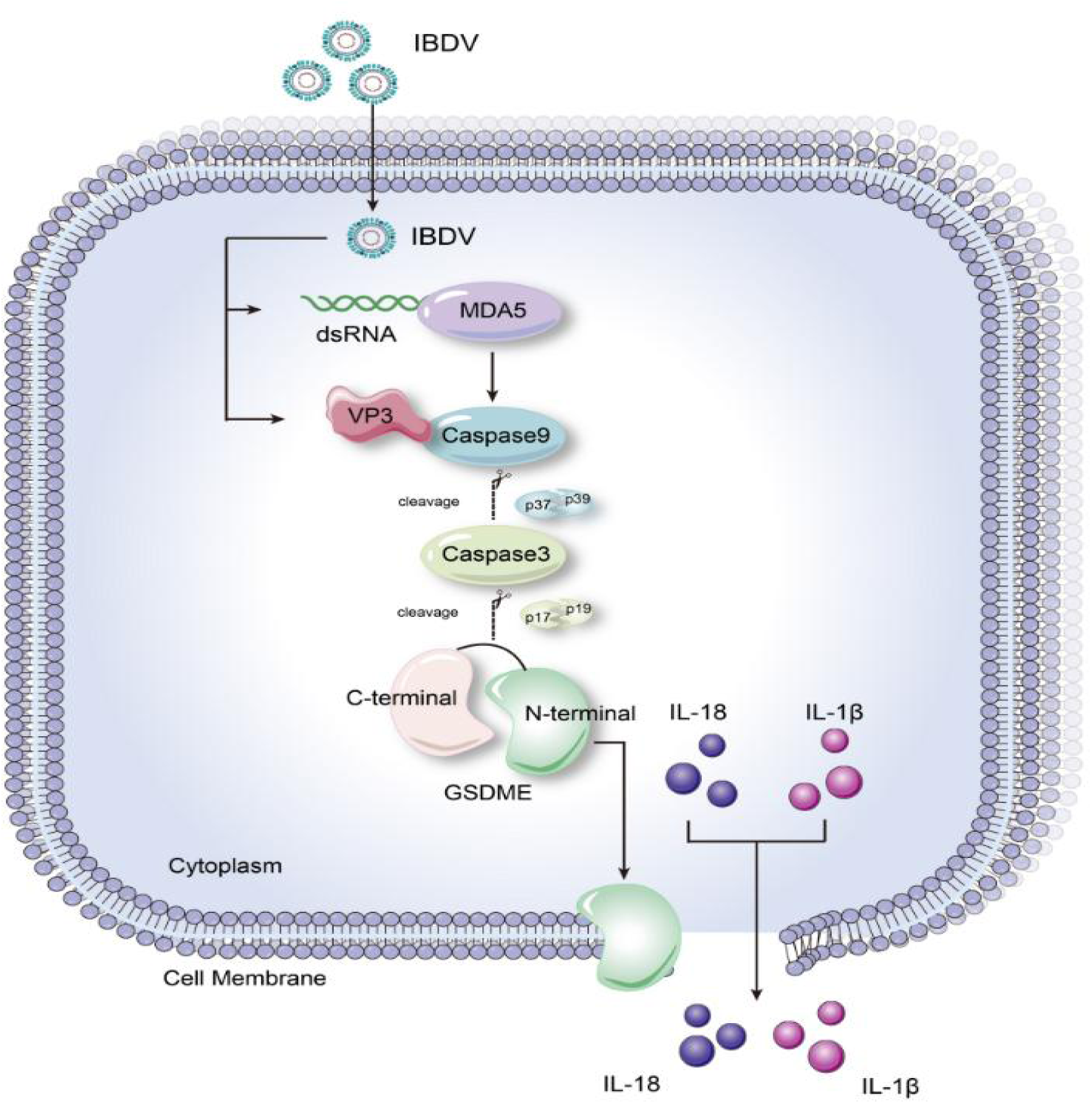
Model of pyroptosis and inflammation induced by nVarIBDV targeting Caspase-9. This schematic diagram illustrates the molecular pathway by which VP3 targets Caspase-9 to trigger pyroptosis of B lymphocyte. VP3 directly binds to and activates Caspase-9. Activated Caspase-9 subsequently activates Caspase-3. Then, Caspase-3 mediates the cleavage of GSDME, generating its active N-terminal fragment that induces pyroptosis and trigger an inflammatory responses. The Caspase-9-Caspase-3-GSDME axis represents the central pathway through which nVarIBDV induces pyroptosis of B lymphocyte.

Inflammation is a common and key pathological marker of bursa of Fabricius damage, typically manifested as infiltration of inflammatory cells including macrophages (24, 36, 37). nVarIBDV infection can cause rapid atrophy of the bursa of Fabricius in chickens and trigger severe inflammatory responses (13). Together, they lead to severe immunosuppression and secondary infections. Previous studies have shown that IBDV infection is accompanied by necrosis and exhaustion of B lymphocyte and significant upregulation of inflammatory factors such as IL-1β, IL-6, IL-8, and IL-12 (16), suggesting that inflammation is closely related to tissue damage. When evaluating the inflammatory responses induced by nVarIBDV, we found that inflammatory cell infiltration and lymphocyte death in the bursa of Fabricius infected with nVarIBDV were observed, and under a scanning electron microscope, the cell membrane of the bursa of Fabricius was broken, similar to the characteristics of pyroptosis. Although the traditional view holds that IBDV mainly induces B lymphocyte apoptosis (38–40) (a non-inflammatory programmed death), this cannot fully explain the intense inflammatory responses during the infection process. Pyroptosis might explain why B lymphocyte exhaustion (atrophy) occurs simultaneously with inflammation during IBDV infection. Although previous studies have reported that attenuated strains of IBDV can induce pyroptosis of chicken fibroblast cell line (DF-1) through the pattern recognition receptor MDA5 (31), it remains unclear whether pathogenic IBDV can induce pyroptosis in B lymphocyte (IBDV target cells) and what the molecular mechanism is. In this study, we confirmed that nVarIBDV induces B lymphocyte pyroptosis by observing the classic "bubbling" morphology in DT40 cells and by measuring the release of LDH and IL-18, thereby showing that it drives inflammatory responses. Therefore, this study provides the first evidence that nVarIBDV infection can trigger an inflammatory responses by inducing pyroptosis of B lymphocyte.

In mammals, virus-induced pyroptosis usually relies on multiple Gasdermin proteins such as GSDMD (40–42), while in poultry, due to the absence of GSDMD, pyroptosis is mainly mediated by GSDMA and GSDME. This study found that nVarIBDV infection could trigger GSDME-dependent pyroptosis, specifically manifested as upregulation of GSDME transcription and protein cleavage, and this process depends on the activation of Caspase-3. Inhibition of Caspase-3 can significantly block the cleavage of GSDME and the release of LDH and IL-18 induced by nVarIBDV infection. Unlike the strategies of Seneca virus (SVV) (43) or foot-and-mouth disease virus (FMDV) (44), which can directly cleft GSDMD/E, nVarIBDV induces pyroptosis and inflammation of B lymphocyte through the conserved Caspase-3-GSDME pathway. It is worth noting that this pathway has also been used by various viruses to induce pyroptosis and inflammation, such as duck hepatitis A virus type 1 (DHAV-1) (45), human enterovirus type 71 (EV71) (46), and H7N9 influenza virus (20). To clarify the specific mechanism by which nVarIBDV activates Caspase-3-GSDME pathway, we screened its viral proteins and found that the structural protein VP3 can effectively promote the activation of Caspase-3 and the cleavage of GSDME. However, Co-IP results indicated that VP3 did not directly bind to Caspase-3, suggesting that its target site was located upstream of Caspase-3. It is known that Caspase-3 can be specifically activated by Caspase-9, which is often used by viruses such as avian encephalomyelitis virus (AEV) (47), equine arteritis virus (EAV) (48), Human Immunodeficiency Virus (HIV) (49) and Japanese Encephalitis Virus (JEV) (50) to initiate apoptosis. While, our study revealed that the nVarIBDV protein VP3 directly binds to and activates Caspase-9 to trigger pyroptosis, with its N-terminal region identified as the key domain for this interaction and for activating Caspase-3-GSDME pathway. Unlike the previous attenuated strain IBDV, which initiates pyroptosis by binding MDA5 to its dsRNA, our results revealed that the viral protein of nVarIBDV can directly initiate the pyroptosis pathway. This mechanism implies that the nVarIBDV viral protein has "reprogrammed" the classic apoptotic initiating molecule Caspase-9, transforming it into a driver of pyroptosis. This strategy is similar to how influenza A virus (IAV) induces pyroptosis by activating Caspase-8 (51, 52). In conclusion, this study not only reveals a new pathway by which nVarIBDV induces inflammatory damage but also expands the known mechanism by which viruses hijack the host apoptotic molecule to induce pyroptosis.

Serine residues often serve as critical sites for stabilizing interactions between viral and host proteins. For instance, phosphorylation of Ser13 in Human cytomegalovirus (HCMV) UL97 promotes binding to 14-3-3 and protein stability (53), while CK2-mediated phosphorylation of Ser23 in human papillomavirus 16 (HPV16) E2 facilitates its interaction with TopBP1 (54). Unlike the above strategies, our study found that the Ser33 site of nVarIBDV VP3 is a key amino acid for initiating Caspase-9, and mutations at this site will significantly weaken the ability of VP3 to activate Caspase-9-Caspase-3-GSDME pathway. Therefore, Ser33 is not only the structural basis for interaction but also the functional core for initiating the downstream pyroptosis pathway. To explore its potential mechanism, we conducted molecular docking and binding energy analyses. These results support a model that Ser33 may help stabilize the VP3-Caspase-9 interaction, possibly through polar contact near the Caspase-9 catalytic region. Consistent with this model, it is predicted that the S33A mutation will reduce interfacial complementarity and increase binding free energy, which is consistent with the observed phenotype of functional loss. Although the precise structural details remain to be further verified, our experimental and computational data consistently emphasize that Ser33 is a key determinant of Caspase-9 activation. In conclusion, this study reveals that VP3 Ser33 is a key target regulating the switching of host cell death during nVarIBDV infection. This provides a direct theoretical basis for a novel antiviral strategy to block pyroptosis and inflammatory damage caused by nVarIBDV.

In summary, we demonstrate that the viral protein VP3 directly activates Caspase-9 to initiate the Caspase-3-GSDME-mediated pyroptosis in B lymphocyte, which underlies the severe inflammatory responses and bursal atrophy characteristic of infectious bursal disease. The identification of Ser33 in VP3 as the critical interface for this interaction not only reveals a key viral determinant of pathogenicity but also pinpoints a precise target for developing therapeutics aimed at disrupting pyroptosis and mitigating the inflammatory damage caused by nVarIBDV.

## Data Availability Statement

The authors confirm that all data underlying the findings are fully available without restriction. All relevant data are within the paper and its supporting information files.

## Acknowledgements

This work was supported by grants from the National Key Research and Development Program of China (2023YFE0106100 and 2022YFD1800300 ), National Natural Science Foundation of China (32473000), Heilongjiang Provincial Natural Science Foundation of China (YQ2023C029), China Agriculture Research System (CARS-41) and Innovation program of Chinese academy of agricultural science (CAAS-CSLPDCP-202402).

Conceptualization:T.Z. and Y.G.; Methodology: T.Z. and S.W.; Formal analysis: T.Z. and Y.G.; Investigation: T.Z., L.T., Y. L., G.W., H.Y., Y. Z., R.Z., N.G., Y.W., W.F., T.C., and Y. Z.; Writing-Original Draft: T.Z. and Y.G.; Writing-Review and Editing: all authors; Visualization: T.Z., X. Q., and Y.G.; Supervision:T.Z. and Y.G.; Project administration: Y.L. and X.Q.; Funding acquisition: Y.G. and S.W.

## References

1. Sharma JM, Kim IJ, Rautenschlein S, Yeh HY. 2000. Infectious bursal disease virus of chickens: pathogenesis and immunosuppression. Dev Comp Immunol 24:223–35.

2. Rautenschlein S, Alkie TN. 2016. Infectious bursal disease virus in poultry: current status and future prospects. Veterinary Medicine: Research and Reports doi:10.2147/vmrr.S689059-18.

3. Mahgoub HA, Bailey M, Kaiser P. 2012. An overview of infectious bursal disease. Arch Virol 157:2047–57.

4. Zierenberg K, Raue R, Nieper H, Islam MR, Eterradossi N, Toquin D, Müller H. 2004. Generation of serotype 1/serotype 2 reassortant viruses of the infectious bursal disease virus and their investigation in vitro and in vivo. Virus Research 105:23–34.

5. Gao H, Wang Y, Gao L, Zheng SJ. 2023. Genetic Insight into the Interaction of IBDV with Host—A Clue to the Development of Novel IBDV Vaccines. International Journal of Molecular Sciences 24:8255.

6. Mahgoub HA. 2012. An overview of infectious bursal disease. Archives of Virology 157:2047–2057.

7. Zhang W, Wang X, Gao Y, Qi X. 2022. The Over-40-Years-Epidemic of Infectious Bursal Disease Virus in China. Viruses 14:2253.

8. Takahashi M, Oguro S, Kato A, Ito S, Tsutsumi N. 2024. Novel Antigenic Variant Infectious Bursal Disease Virus Outbreaks in Japan from 2014 to 2023 and Characterization of an Isolate from Chicken. Pathogens 13:1141.

9. Thai TN, Jang I, Kim H-A, Kim H-S, Kwon Y-K, Kim H-R. 2021. Characterization of antigenic variant infectious bursal disease virus strains identified in South Korea. Avian Pathology 50:174–181.

10. Aliyu HB, Hair-Bejo M, Omar AR, Ideris A. 2021. Genetic Diversity of Recent Infectious Bursal Disease Viruses Isolated From Vaccinated Poultry Flocks in Malaysia. Front Vet Sci 8:643976.

11. Jaton J, Lozano LC, Gambini P, Ponti M, GцЁmez E, KцІnig GA, Chimeno Zoth S. 2024. Research Note: Characterization and phylodynamic analysis of new infectious bursal disease virus variants circulating in Argentina. Poult Sci 103:103623.

12. Shahein MA, Sultan HA, Zanaty A, Adel A, Mosaad Z, Said D, Erfan A, Samy M, Selim A, Selim K, Naguib MM, Hassan H, Shazly OE, El-Badiea ZA, Moawad MK, Samir A, Shahaby ME, Farghaly E, Eid S, Abdelaziz MN, Hamoud MM, Mehana O, Hagag NM, Samy A. 2024. Emergence of the novel infectious bursal disease virus variant in vaccinated poultry flocks in Egypt. Avian Pathol 53:419–429.

13. Fan L, Wu T, Wang Y, Hussain A, Jiang N, Gao L, Li K, Gao Y, Liu C, Cui H, Pan Q, Zhang Y, Wang X, Qi X. 2020. Novel variants of infectious bursal disease virus can severely damage the bursa of fabricius of immunized chickens. Veterinary Microbiology 240:108507.

14. Li K, Niu X, Jiang N, Zhang W, Wang G, Li K, Huang M, Gao Y, Qi X, Wang X. 2023. Comparative Pathogenicity of Three Strains of Infectious Bursal Disease Virus Closely Related to Poultry Industry. Viruses 15:1257.

15. Spackman E, Stephens CB, Pantin-Jackwood MJ. 2018. The Effect of Infectious Bursal Disease Virus-Induced Immunosuppression on Vaccination Against Highly Pathogenic Avian Influenza Virus. Avian Dis 62:36–44.

16. Fan L, Wang Y, Jiang N, Chen M, Gao L, Li K, Gao Y, Cui H, Pan Q, Liu C, Zhang Y, Wang X, Qi X. 2020. Novel variant infectious bursal disease virus suppresses Newcastle disease vaccination in broiler and layer chickens. Poultry Science 99:6542–6548.

17. Laghlali G, Lawlor KE, Tate MD. 2020. Die Another Way: Interplay between Influenza A Virus, Inflammation and Cell Death. Viruses 12:401.

18. Li S, Zhang Y, Guan Z, Ye M, Li H, You M, Zhou Z, Zhang C, Zhang F, Lu B, Zhou P, Peng K. 2023. SARS-CoV-2 Z-RNA activates the ZBP1-RIPK3 pathway to promote virus-induced inflammatory responses. Cell Research 33:201–214.

19. Kalejta RF, Zhang X, Chen G, Yin J, Nie L, Li L, Du Q, Tong D, Huang Y. 2024. Pseudorabies Virus UL4 protein promotes the ASC-dependent inflammasome activation and pyroptosis to exacerbate inflammation. PLOS Pathogens 20:e1012546.

20. Wan X, Li J, Wang Y, Yu X, He X, Shi J, Deng G, Zeng X, Tian G, Li Y, Jiang Y, Guan Y, Li C, Shao F, Chen H. 2022. H7N9 virus infection triggers lethal cytokine storm by activating gasdermin E-mediated pyroptosis of lung alveolar epithelial cells. National Science Review 9:

21. Feng Y, Li M, Yangzhong X, Zhang X, Zu A, Hou Y, Li L, Sun S. 2022. Pyroptosis in inflammation-related respiratory disease. Journal of Physiology and Biochemistry 78:721–737.

22. Min R, Bai Y, Wang NR, Liu X. 2025. Gasdermins in pyroptosis, inflammation, and cancer. Trends Mol Med 31:860–875.

23. Li M, Zhu Y, Li M. 2025. Newcastle Disease Virus Induces Pyroptosis in Canine Mammary Tumour CMT-U27 Cells via the TNFα/NF-κB/NLRP3 Signalling Pathway. Vet Comp Oncol 23:224–235.

24. Asfor AS, Nazki S, Reddy VRAP, Campbell E, Dulwich KL, Giotis ES, Skinner MA, Broadbent AJ. 2021. Transcriptomic Analysis of Inbred Chicken Lines Reveals Infectious Bursal Disease Severity Is Associated with Greater Bursal Inflammation In Vivo and More Rapid Induction of Pro-Inflammatory Responses in Primary Bursal Cells Stimulated Ex Vivo. Viruses 13:933.

25. Huang M, Xu M, Han J, Ke E, Niu X, Zhang Y, Wang G, Yu H, Liu R, Wang S, Liu Y, Chen Y, Han J, Wu Z, Cui H, Zhang Y, Duan Y, Gao Y, Qi X. 2025. Enhancing MyD88 oligomerization is one important mechanism by which IBDV VP2 induces inflammatory response. PLoS Pathog 21:e1012985.

26. Fan L, Wu T, Hussain A, Gao Y, Zeng X, Wang Y, Gao L, Li K, Wang Y, Liu C, Cui H, Pan Q, Zhang Y, Liu Y, He H, Wang X, Qi X. 2019. Novel variant strains of infectious bursal disease virus isolated in China. Vet Microbiol 230:212–220.

27. Percie du Sert N, Hurst V, Ahluwalia A, Alam S, Avey MT, Baker M, Browne WJ, Clark A, Cuthill IC, Dirnagl U, Emerson M, Garner P, Holgate ST, Howells DW, Karp NA, Lazic SE, Lidster K, MacCallum CJ, Macleod M, Pearl EJ, Petersen OH, Rawle F, Reynolds P, Rooney K, Sena ES, Silberberg SD, Steckler T, Wц╪rbel H. 2020. The ARRIVE guidelines 2.0: Updated guidelines for reporting animal research. PLoS Biol 18:e3000410.

28. Adasme MF, Linnemann KL, Bolz SN, Kaiser F, Salentin S, Haupt VJ, Schroeder M. 2021. PLIP 2021: expanding the scope of the protein-ligand interaction profiler to DNA and RNA. Nucleic Acids Res 49:W530–w534.

29. Laskowski RA, Swindells MB. 2011. LigPlot+: multiple ligand-protein interaction diagrams for drug discovery. J Chem Inf Model 51:2778–86.

30. Liu J, Wang X, Wang X, Wang J, Ma Y, Cao Y, Zhang W. 2024. Chicken gasdermins mediate pyroptosis after the cleavage by caspases. International Journal of Biological Macromolecules 270:132476.

31. Chen Z, Chang H, Zhang S, Gao H, Gao L, Cao H, Li X, Wang Y, Zheng SJ. 2025. Chicken GSDME, a major pore-forming molecule responsible for RNA virus-induced pyroptosis in chicken. J Virol 99:e0158824.

32. Bhat AA, Thapa R, Afzal O, Agrawal N, Almalki WH, Kazmi I, Alzarea SI, Altamimi ASA, Prasher P, Singh SK, Dua K, Gupta G. 2023. The pyroptotic role of Caspase-3/GSDME signalling pathway among various cancer: A Review. International Journal of Biological Macromolecules 242:124832.

33. Yin Q, Park HH, Chung JY, Lin SC, Lo YC, da Graca LS, Jiang X, Wu H. 2006. Caspase-9 holoenzyme is a specific and optimal procaspase-3 processing machine. Mol Cell 22:259–68.

34. Yu H, Wang G, Zhang W, Wu Z, Niu X, Huang M, Zhang Y, Liu R, Han J, Xu M, Han J, Ling D, Ke E, Wang S, Cui H, Zhang Y, Chen Y, Liu Y, Duan Y, Gao Y, Qi X. 2025. Epidemiological characteristics of infectious bursal disease virus (IBDV) in China from 2023 to 2024: Mutated very virulent IBDV (mvvIBDV) is associated with atypical IBD. Poult Sci 104:105195.

35. Jiang N, Wang Y, Zhang W, Niu X, Huang M, Gao Y, Liu A, Gao L, Li K, Pan Q, Liu C, Zhang Y, Cui H, Wang X, Qi X. 2021. Genotyping and Molecular Characterization of Infectious Bursal Disease Virus Identified in Important Poultry-Raising Areas of China During 2019 and 2020. Front Vet Sci 8:759861.

36. Yu Y, Xu Z, Zhang Y, Wang Q, Ou C, Wang Y, Wang L, Gao P, Du S, Guo F, Ma J. 2020. Ghrelin attenuates infectious bursal disease virus–induced early inflammatory response and bursal injury in chicken. Poultry Science 99:5399–5406.

37. Xu Z, Yu Y, Liu Y, Ou C, Zhang Y, Liu T, Wang Q, Ma J. 2019. Differential expression of pro-inflammatory and anti-inflammatory genes of layer chicken bursa after experimental infection with infectious bursal disease virus. Poultry Science 98:5307–5314.

38. Qin Y, Xu Z, Wang Y, Li X, Cao H, Zheng SJ. 2017. VP2 of Infectious Bursal Disease Virus Induces Apoptosis via Triggering Oral Cancer Overexpressed 1 (ORAOV1) Protein Degradation. Frontiers in Microbiology 8:1351.

39. Rodriguez-Lecompte JC, Nino-Fong R, Lopez A, Frederick Markham RJ, Kibenge FS. 2005. Infectious bursal disease virus (IBDV) induces apoptosis in chicken B cells. Comp Immunol Microbiol Infect Dis 28:321–37.

40. Duan X, Zhao M, Wang Y, Li X, Cao H, Zheng SJ. 2020. Epigenetic Upregulation of Chicken MicroRNA-16-5p Expression in DF-1 Cells following Infection with Infectious Bursal Disease Virus (IBDV) Enhances IBDV-Induced Apoptosis and Viral Replication. J Virol 94:e01724–19.

41. Man SM, Karki R, Kanneganti TD. 2017. Molecular mechanisms and functions of pyroptosis, inflammatory caspases and inflammasomes in infectious diseases. Immunol Rev 277:61–75.

42. Lin R, Porto BN. 2025. Pyroptosis in Respiratory Virus Infections: A Narrative Review of Mechanisms, Pathophysiology, and Potential Therapeutic Interventions. Microorganisms 13:2109.

43. Wen W, Li X, Wang H, Zhao Q, Yin M, Liu W, Chen H, Qian P. 2021. Seneca Valley Virus 3C Protease Induces Pyroptosis by Directly Cleaving Porcine Gasdermin D. The Journal of Immunology 207:189–199.

44. Ren X, Yin M, Zhao Q, Zheng Z, Wang H, Lu Z, Li X, Qian P. 2023. Foot-and-Mouth Disease Virus Induces Porcine Gasdermin E-Mediated Pyroptosis through the Protease Activity of 3C(pro). J Virol 97:e0068623.

45. Wang J, Yan H, Bei L, Jiang S, Zhang R. 2024. 2A2 protein of DHAV-1 induces duck embryo fibroblasts gasdermin E-mediated pyroptosis. Veterinary Microbiology 290:109987.

46. Liu T, Wang B, Li Y, Tian S, Gao X, Fan Y, Zhang X, Yang L, Wu R, Liu L. 2025. Enterovirus 71 infection induces pyroptotic brain injury via synergistic activation of classical inflammasome and viral gasdermin D cleavage. J Virol doi:10.1128/jvi.01860-25e0186025.

47. Liu J, Wei T, Kwang J. 2004. Avian encephalomyelitis virus nonstructural protein 2C induces apoptosis by activating cytochrome c/caspase-9 pathway. Virology 318:169–182.

48. St-Louis M-C, Archambault D. 2007. The equine arteritis virus induces apoptosis via caspase-8 and mitochondria-dependent caspase-9 activation. Virology 367:147–155.

49. Muthumani K, Hwang DS, Desai BM, Zhang D, Dayes N, Green DR, Weiner DB. 2002. HIV-1 Vpr Induces Apoptosis through Caspase 9 in T Cells and Peripheral Blood Mononuclear Cells. Journal of Biological Chemistry 277:37820–37831.

50. Tsao C-H, Su H-L, Lin Y-L, Yu H-P, Kuo S-M, Shen C-I, Chen C-W, Liao C-L. 2008. Japanese encephalitis virus infection activates caspase-8 and -9 in a FADD-independent and mitochondrion-dependent manner. Journal of General Virology 89:1930–1941.

51. Sun Y, Yu H, Zhan Z, Liu W, Liu P, Sun J, Zhang P, Wang X, Liu X, Xu X. 2025. TRIF-TAK1 signaling suppresses caspase-8/3-mediated GSDMD/E activation and pyroptosis in influenza A virus-infected airway epithelial cells. iScience 28:111581.

52. Kuriakose T, Man SM, Subbarao Malireddi RK, Karki R, Kesavardhana S, Place DE, Neale G, Vogel P, Kanneganti T-D. 2016. ZBP1/DAI is an innate sensor of influenza virus triggering the NLRP3 inflammasome and programmed cell death pathways. Science Immunology 1:aag2045.

53. Iwahori S, Umaña AC, Kalejta RF, Murata T. 2022. Serine 13 of the human cytomegalovirus viral cyclin-dependent kinase UL97 is required for regulatory protein 14-3-3 binding and UL97 stability. Journal of Biological Chemistry 298:102513.

54. Prabhakar AT, James CD, Das D, Otoa R, Day M, Burgner J, Fontan CT, Wang X, Glass SH, Wieland A, Donaldson MM, Bristol ML, Li R, Oliver AW, Pearl LH, Smith BO, Morgan IM, Laimins LA. 2021. CK2 Phosphorylation of Human Papillomavirus 16 E2 on Serine 23 Promotes Interaction with TopBP1 and Is Critical for E2 Interaction with Mitotic Chromatin and the Viral Life Cycle. mBio 12:e0116321.

